# Assembly and seasonality of core phyllosphere microbiota on perennial biofuel crops

**DOI:** 10.1101/446369

**Authors:** Keara L Grady, Jackson W. Sorensen, Nejc Stopnisek, John Guittar, Ashley Shade

**Author notes:** Contributed equally (Keara L Grady, Jackson W. Sorensen, Nejc Stopnisek).

## Abstract

Perennial grasses are promising feedstocks for biofuel production, and there is potential to leverage their native microbiomes to increase their productivity and resilience to environmental stress. Here, we characterize the 16S rRNA gene diversity and seasonal assembly of bacterial and archaeal microbiomes of two perennial cellulosic feedstocks, switchgrass (Panicum virgatum L.) and miscanthus (Miscanthus × giganteus). We sampled leaves and soil every three weeks from pre-emergence through senescence for two consecutive switchgrass growing seasons and one miscanthus season, and identified core leaf taxa based on abundance and occupancy. Virtually all leaf taxa are also detected in soil; source-sink modeling shows non-random, ecological filtering by the leaf, suggesting that soil is important reservoir of phyllosphere diversity. Core leaf taxa include early, mid, and late season groups that were consistent across years and crops. This consistency in leaf microbiome dynamics and core members is promising for microbiome manipulation or management to support biofuel crop production.

---

The phyllosphere (aerial parts of plants) represents the largest environmental surface area of microbial habitation on the planet ^1–3^, and much of that surface area is cultivated agriculture, including an estimated 1.5 × 10^7^ km^2^ of cropland ^4^. Phyllosphere microorganisms may provide numerous benefits to plants, including increased stress tolerance ^5–7^, promotion of growth and reproduction ^8–10^, protection from foliar pathogens ^11^, and, with soil microbes, control of flowering phenology ^12^. Phyllosphere microorganisms are also thought to play important roles in Earth’s biogeochemical cycles by moderating methanol emissions from plants ^13,14^ and contributing to global nitrogen fixation ^15^. Despite this importance, knowledge of phyllosphere microbiomes remains relatively modest, especially for agricultural crops ^3,16–18^. To leverage plant microbiomes to support productivity and resilience both above and below ground ^19–21^, there is a need to advance foundational knowledge of phyllosphere microbiome diversity and dynamics.

Biofuel crops like miscanthus and switchgrass are selected to have extended growing seasons, to produce ample phyllosphere biomass, and to maintain high productivity when grown on marginal lands that are not optimal for food agriculture ^22–25^. In the field, these grasses provide extensive leaf habitat, with a seasonal maximum leaf area index (LAI) of 6.2 for switchgrass, and 10 m^2^ leaf surface per m^2^ land for miscanthus ^22^, as compared to a maximum LAI of 3.2 for corn ^26^. Upon senescence, the aboveground biomass is harvested for conversion to biofuels and related bioproducts. Improved understanding of the phyllosphere microbiome is expected to advance goals to predict or manage changes in biomass quality in response to abiotic stress like drought ^27–31^ or biotic stress like foliar pathogens ^32–34^.

Leveraging the Great Lakes Bioenergy Research Center’s Biofuel Cropping System Experiment (BCSE; a randomized block design established at Michigan State’s Kellogg Biological Station in 2008), we asked two questions of the bacterial and archaeal communities (henceforth: “microbiomes”) inhabiting the leaf surfaces and the associated soils of switchgrass and miscanthus: 1) Are there seasonal patterns of phyllosphere microbiome assembly? If so, are these patterns consistent across fields of the same crop, different crops, and years? 2) To what extent might soil serve as a reservoir of phyllosphere diversity?

## Results and Discussion

### Sequencing summary and alpha diversity

In total, we sequenced 373 phyllosphere epiphyte (leaf surface) and soil samples across the two growing seasons in 2016 and 2017. The number of sequences per sample after our 97% OTU (operational taxonomic unit) clustering pipeline ranged from 20,647 to 359,553. The percentage of sequences belonging to chloroplasts and mitochondria per sample range between 0.2-99.8%, but 235 of the samples (63%) had fewer than 10% chloroplasts and mitochondria reads. After removing sequences that were attributed to chloroplasts and mitochondria or that had unassigned taxonomic classification, we filtered samples that contained fewer than 1000 reads and rarefied the remaining samples to 1000 reads for comparative analyses. While this number of reads is not sufficient to fully capture soil diversity, it does capture phyllosphere diversity (**Figure 1A**). The majority of the switchgrass and miscanthus phyllosphere communities were exhaustively sequenced, and approached richness asymptotes with their associated soils.

**Figure 1.**
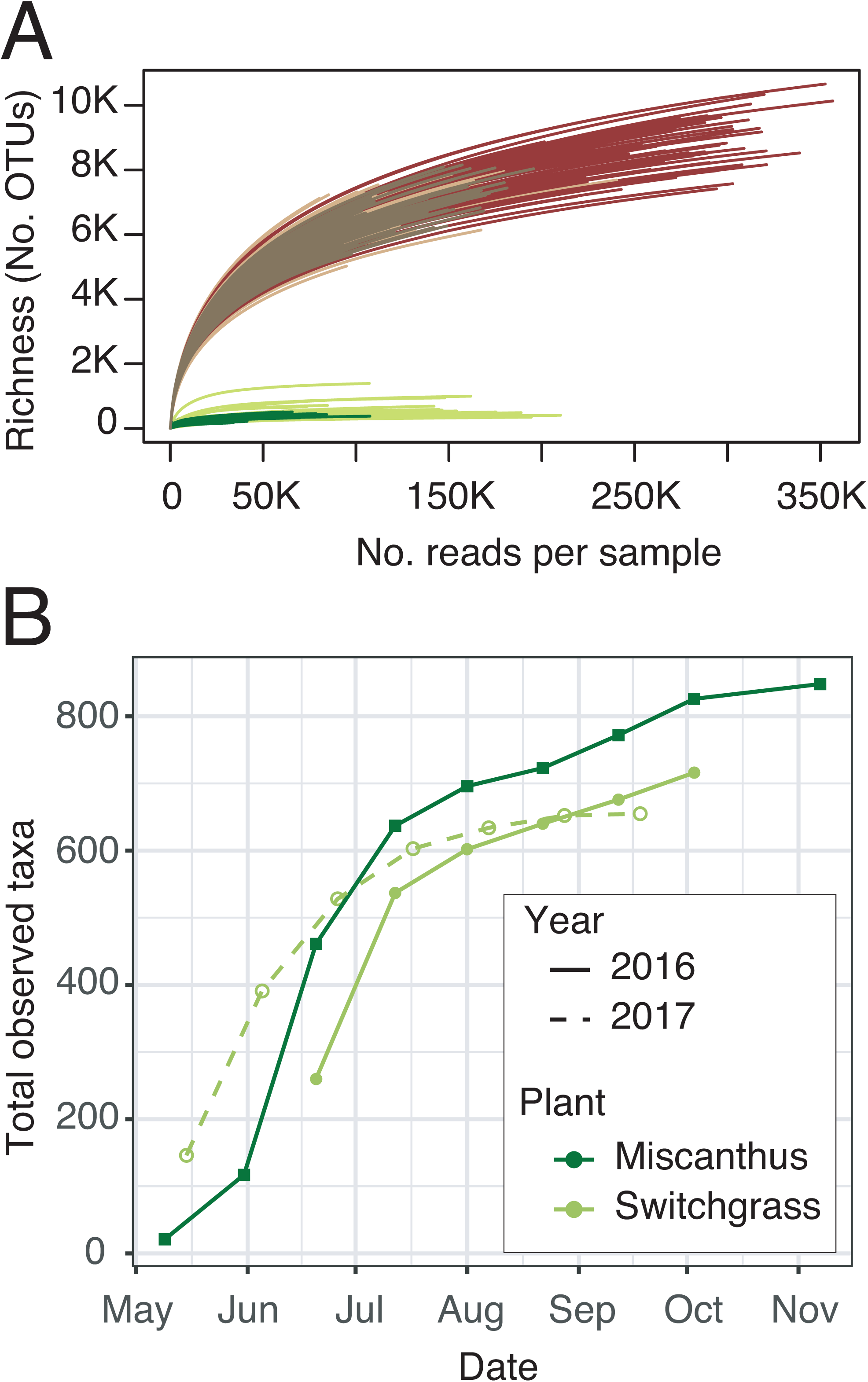
Sequencing effort and alpha diversity for switchgrass and miscanthus phyllosphere and soils. Operational taxonomic units (OTUs) were defined at 97% sequence identity of 16S rRNA gene amplicons. (A) Rarefaction curves of quality-controlled reads. The vertical line is the maximum number of sequences observed in a sample, and the horizontal line is the richness of that sample. (B) Phyllosphere richness accumulation over time, using a dataset subsampled to 1000 sequences per sample.

As reported for other plants ^3,35^, switchgrass and miscanthus phyllosphere communities had relatively low richness, with 1480 total taxa observed across both crops and consistently fewer than 150 taxa per time point, though there was modest seasonal variability in richness (**Figure S1**). Cumulative richness increased most between the two earliest time points, and then tapered gradually upward until senescence (**Figure 1B**), showing that the contributions of new taxa to community richness were low but consistent over time.

### Seasonal microbiome dynamics

To perform the most complete temporal analyses of phyllosphere microbiome seasonality, we also subsampled the amplicon sequencing dataset to include the maximum number of time points, resulting in inclusion of 51 discrete leaf and soil samples collected over 18 total time points. The overarching patterns in beta diversity were consistent and statistically indistinguishable from those derived from the same dataset to include more reads per sample but fewer time points (Mantel tests all p < 0.001; **Table S1, Table S2**, **Figure S2**). For consistency, we report the patterns from 1000 reads per sample in the main text, but for transparency and comparison, we report results from the minimum reads per sample, inclusive of the complete time series, in supporting materials.

There were directional seasonal changes in the structures of switchgrass and miscanthus phyllosphere bacterial and archaeal communities (**Figure 2A**, **Table 1**), and these could be attributed to changes in both soil and leaf properties, as well as to weather (**Table S3**). Over the 2016 season, miscanthus and switchgrass phyllosphere communities were synchronous (changed at the same pace and to the same extent, Procrustes m12= 0.349, R = 0.807, p = 0.021), and community structure became less variable as the growing season progressed (**Figure S3**). Switchgrass 2016 and 2017 leaf communities were highly synchronous, suggesting a predictable, interannual assembly (Procrustes m12= 0.011, R= 0.994, p = 0.008). The switchgrass community structures were overall equivalent between 2016 and 2017, with the exception of the final time points that were collected post-senescence. Together with the species accumulation analysis (**Figure 1B**), these data suggest that these phyllosphere communities are not stochastically assembled, nor are they a linear accumulation over seasonal leaf exposure to whatever taxa are dispersed. The communities follow a directional assembly over the growing season, and the assembly was highly consistent over two years in the switchgrass.

**Figure 2.**
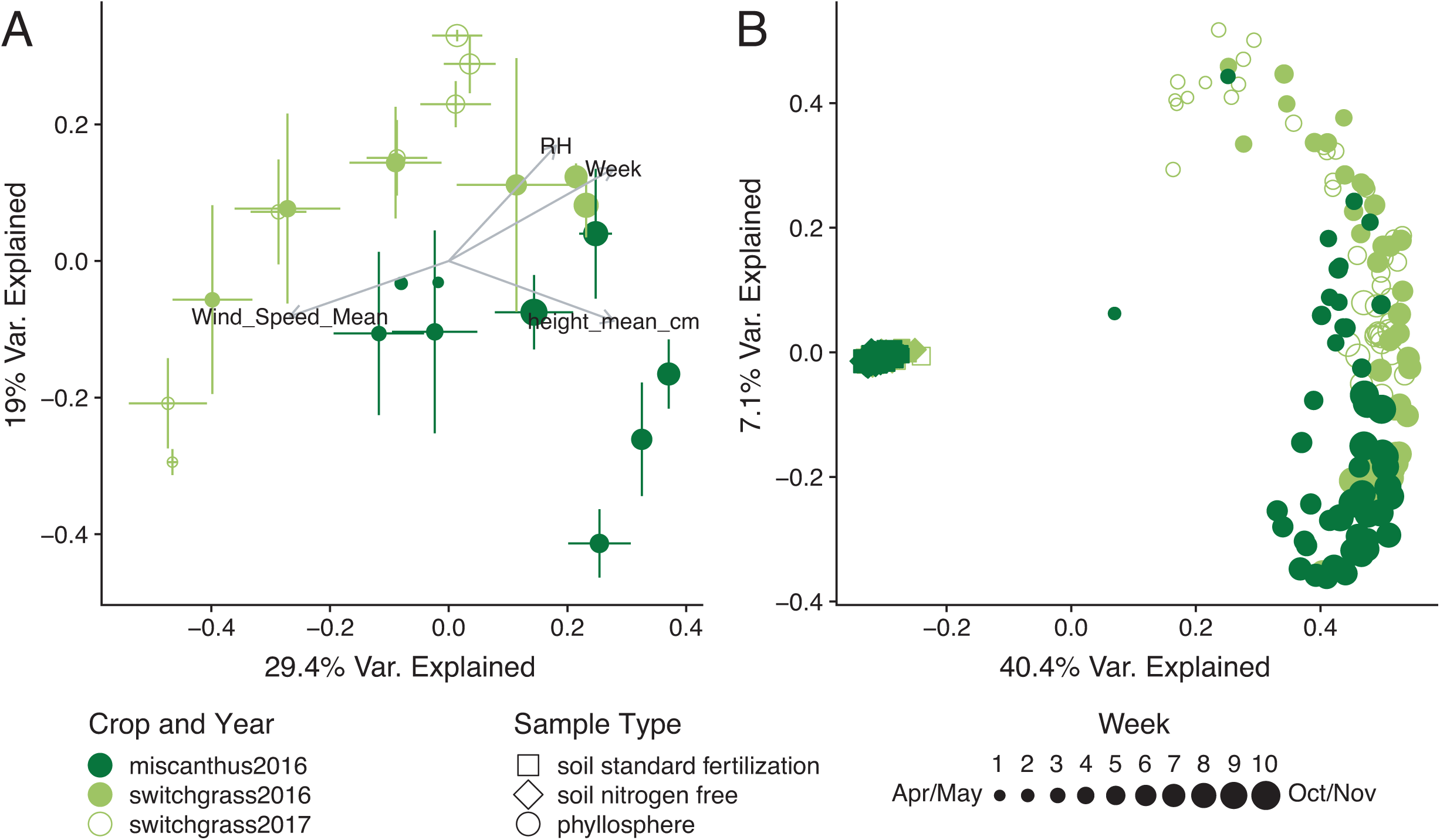
Seasonal patterns in the structures of bacterial and archaeal communities inhabiting the phyllosphere and associated soils of the biofuel feedstocks switchgrass and miscanthus. (A) Principal coordinates analysis (PCoA) of switchgrass and miscanthus phyllosphere communities (Bray-Curtis dissimilarity), error bars show 1 deviation around the centroid (n = 1 to 8 replicate plots/time point). (B) PCoA of the phyllosphere communities relative to the soil. For both A and B, subsampling depth was 1000 reads per sample and environmental vectors were fitted when r^2^ > 0.4 and p < 0.05.

**Table 1.** Permuted multivariate analysis of variance (PERMANOVA) tables for all hypothesis tests for differences in community structure (beta diversity).

| Dataset | Variable Tested | Degrees of Freedom | PseudoF | R squared | p-value |
| --- | --- | --- | --- | --- | --- |
| 2016 All | Microbiome habitat (soil v. leaf) | 1 | 166.09 | 0.415 | 0.001 |
| 2017 All | Microbiome habitat (soil v. leaf) | 1 | 74.66 | 0.418 | 0.001 |
| 2016 Phyllosphere | Time | 1 | 21.02 | 0.183 | 0.001 |
| 2016 Soil | Time | 1 | 7.31 | 0.050 | 0.001 |
| 2017 Phyllosphere | Time | 1 | 29.09 | 0.409 | 0.001 |
| 2017 Soil | Time | 1 | 3.05 | 0.048 | 0.001 |
| 2016 Phyllosphere | Crop | 1 | 14.50 | 0.134 | 0.001 |
| 2016 Soil | Crop | 1 | 6.76 | 0.047 | 0.001 |
| 2016 Phyllosphere | Fertilization status | 1 | 0.40 | 0.004 | 0.95 |
| 2016 Soil | Fertilization status | 1 | 4.77 | 0.033 | 0.001 |
| 2017 Phyllosphere | Fertilization status | 1 | 0.87 | 0.020 | 0.438 |
| 2017 Soil | Fertilization status | 1 | 3.41 | 0.054 | 0.001 |

### Contribution of soil microorganisms to phyllosphere assembly

The major sources of microorganisms to the phyllosphere are soils ^2^, the vascular tissue of the plant or its seed ^36^, and the atmosphere or arthropod vectors ^3^. As several studies have shown that soil microbes contribute to the phyllosphere microbiome ^35,37^, we wanted to understand the potential for soil as a reservoir of microorganisms inhabiting switchgrass and miscanthus phyllospheres. We hypothesized that the intersect of shared soil and phyllosphere taxa would be highest early in the season, after the young grasses emerged through the soil. Our deep sequencing effort also provided the opportunity to investigate differences in taxon relative abundances between soil and leaf communities, and to understand what contributions, if any, the soil rare biosphere has for leaf assembly.

First, we interrogated the 2016 time series to determine the influence of soil-detected taxa on leaf microbial communities for both crops. As expected, the structures of leaf communities were highly distinct from soils (**Figure 2B**, **Figure S2B**, **Table 1**). Though soil communities also changed seasonally, they experienced less overall change than the phyllosphere (**Table 1**, **Figure 2B**, **Figure S2C**). While fertilization had no impact on phyllosphere communities, it did have small but significant influence on soil communities (**Table 1**). These seasonal and fertilization treatment patterns were reproduced in 2017 for switchgrass (**Table 1**).

To better understand the relationship between soil and phyllosphere communities, including the influence of rare members of the soil, we searched for phyllosphere taxa within the full soil dataset (not subsampled). Approximately 90% of phyllosphere OTUs were present in soil samples, with negligible differences between the two crops and modest variability over time (**Figure 3A**). When considering the relative abundances of taxa, the mean fraction of phyllosphere communities found in soil samples was even higher at 98% and exhibited no clear trend over time (**Figure 3B**). Our results show that the majority of abundant, commonly detected taxa in the phyllosphere are also present in the soil – albeit often at very low abundances (see below) – highlighting a potentially important role of the soil in harboring phyllosphere taxa between plant colonization events.

**Figure 3.**
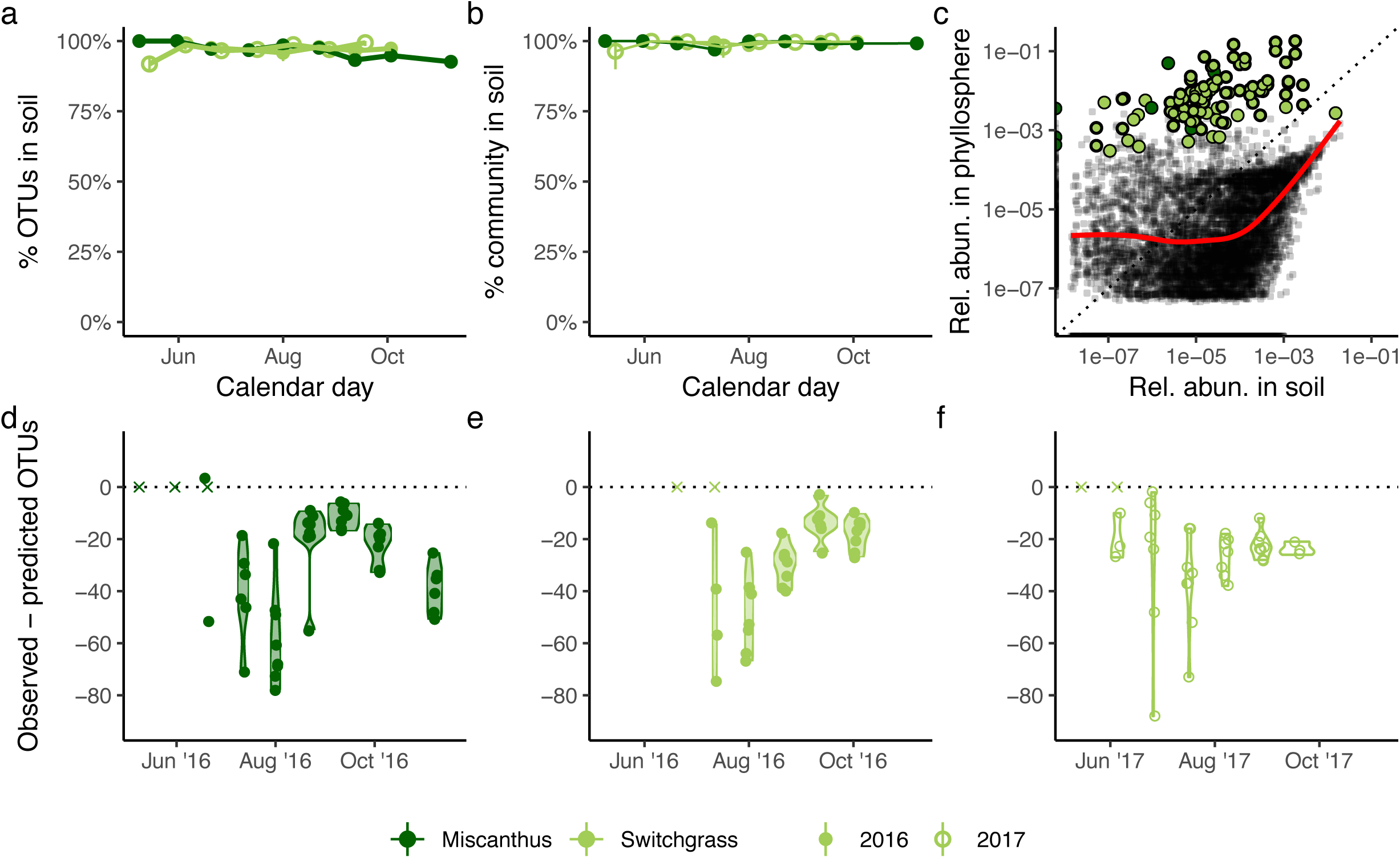
The majority of phyllosphere taxa were also present in the soil. (**A**) Circles represent the mean number of OTUs found in up to eight replicate phyllosphere samples, subsampled to 1000 sequences, for each crop at each time point. An OTU was considered present in the soil if it occurred at any abundance in any of the 202 unrarefied soil samples over two years. **(B)** The fractions of the phyllosphere communities present in the soil were even greater when considering the relative abundances of taxa; each circle represents the mean total relative abundance of leaf taxa present in the soil in up to four replicate phyllosphere samples. **(C)** The relative abundances of taxa in pooled phyllosphere samples and pooled soil samples were positively correlated among taxa that were present at greater than 0.01% total abundance in the soil. Each black circle represents an OTU present in both phyllosphere and soil communities; a LOESS smoothing function is shown as a red line. Core members are shown as large green circles. (**D-F**) Source-sink models of phyllosphere community assembly from soils. Violin plots show the numbers of observed taxa in the phyllosphere were consistently lower than the richness values predicted by model simulations. The model assumed random increases and decreases in taxon abundances between time points and random immigration from the soil community (see Methods). Each circle represents a single phyllosphere sample.

Given the large proportion of phyllosphere taxa present in the soil, we explored the potential role that immigration from the soil may have in shaping phyllosphere community composition and seasonal patterns. On balance, many abundant and persistent phyllosphere taxa were in low abundances in the soil, though there was a positive association between soil and leaf abundances for soil taxa > 1 e04 relative abundance (**Figure 3C**). This result indirectly supports the presence of an ecological filter operating on the phyllosphere that favors some taxa while disfavoring others. To further investigate how ecological filtering may vary over the growing season, we compared observed trends in OTU richness with those predicted by a source-sink null model which simulated demographic stochasticity and random immigration from the soil between subsequent sampling points (see Methods, **Figure 3D-F**). Observed phyllosphere communities were dramatically less rich than null model predictions, again supporting the presence of a strong ecological filter. Such a filter could be due to host plant selection, environmental filtering, competition exclusion among microbial taxa, or a combination of all three. According to this model, the strength of filtering did not trend consistently over the growing season.

Finally, other studies have found that soil microbes contribute more to early-season phyllosphere communities ^38^, and we observed similar patterns: the most abundant soil taxa that were also detected on leaves were more prominent in the early season and then became rare and transient on leaves in the late season (**Figure S4**).

We conclude from these results that soil is a major reservoir of leaf microorganisms for these perennial crops and note that deep sequencing was required of the soils to observe many of the prominent leaf taxa. This is in contrast to the studies of other plants that have suggested that the phyllosphere is comprised largely of passively dispersed and stochastically assembled microbes from the atmosphere ^39–41^. Notably, our analysis cannot inform directionality or mechanism of dispersal, which could have occurred between soil and leaf via wind, insects, or through grass emergence, etc. While 133 leaf OTUs (9%) observed in the phyllosphere could not be detected in the soil (using the unrarified soil dataset) and may be attributable to non-soil reservoirs, the vast majority of leaf microbes were detectable in local soils and non-neutral assembly patterns suggest both determinism and habitat filtering.

### Core members of the switchgrass and miscanthus phyllosphere

There was high overlap between switchgrass and miscanthus phyllosphere communities and a trend towards increased intra-crop similarity during senescence (**Figure S5**). There was also a modest influence of host crop in 2016 (**Table 1**). Therefore, we defined a core microbiome for each crop and season (**Dataset 1**). We applied an established macroecological approach ^42^ to consider both the occupancy and abundance patterns of these taxa (**Figure 4A-C**); abundance-occupancy relationships have been previously explored for microbial communities ^43,44^ and we utilize it here for ecologically informing a core microbiome. *Occupancy* is an ecological term that considers how consistently a taxon is detected across samples in the dataset, expressed as a proportion of occurrences given the total samples collected (e.g., 1.0 or 100%). Occupancy provides a dataset-aggregated term describing taxon persistence, which can be informative for defining core taxa when datasets include a time series ^45^.

**Figure 4.**
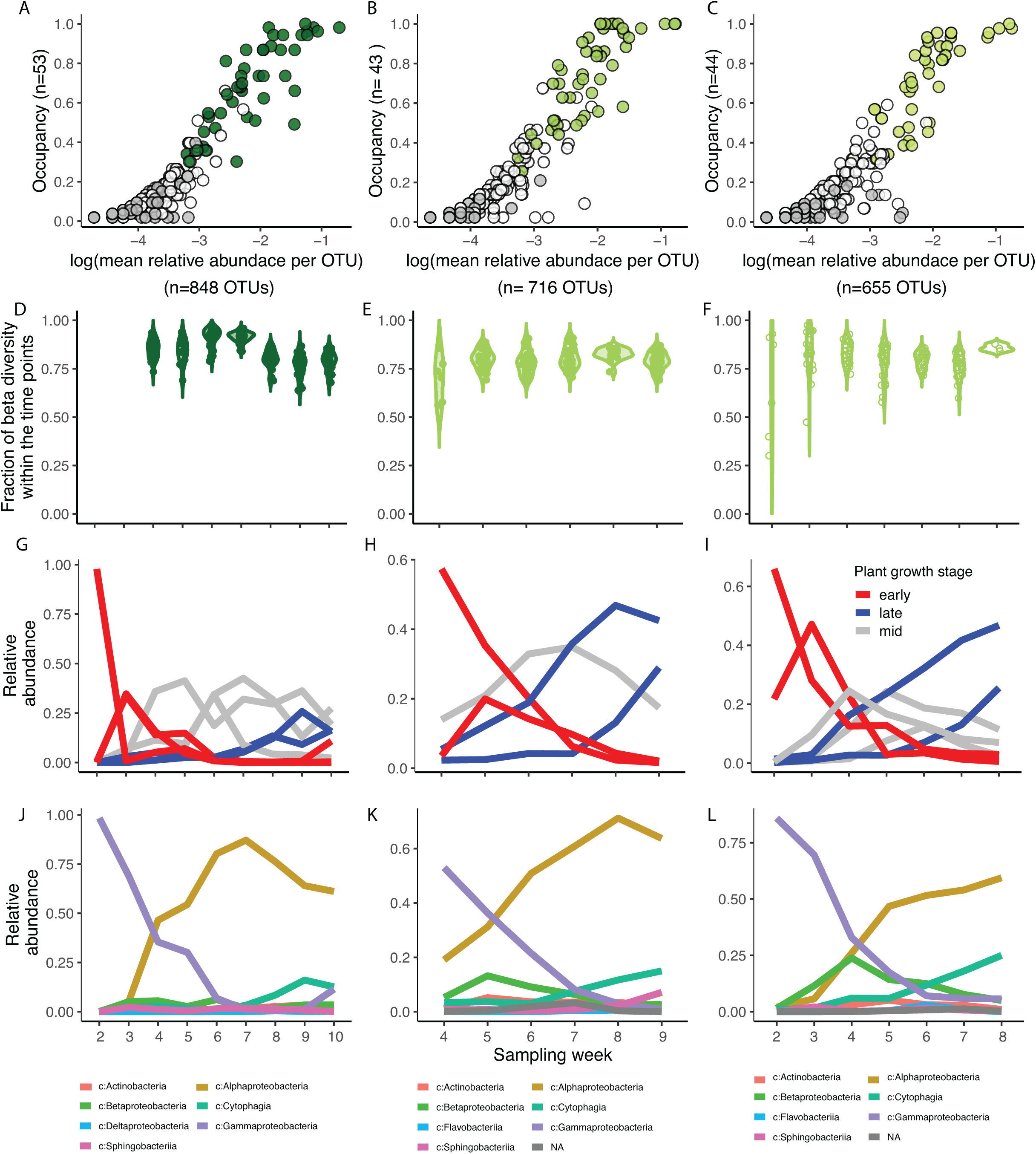
Selection and dynamics of core phyllosphere members. Abundance-occupancy of leaf taxa for **(A)** miscanthus 2016, **(B)** switchgrass 2016, and **(C)** switchgrass 2017, and their inclusion in their respective cores. Each point is an OTU. Abundance-occupancy distributions were calculated at each time point, and taxa that had 100% occupancy at any time point (e.g., were detected in all replicate plots at one sampling date) were included in the core (green). Non-core taxa that were detected in both crops (white/open circles), and crop-specific taxa (grey) are also indicated. (**D-F**) Contributions of the core taxa to changes in beta diversity over time. (**G-I**) Patterns of core taxa that share similar temporal changes, as determined by hierarchical clustering of standardized dynamics. Colors correspond to the dendrograms in **Figure S7**. (**J-L**) Patterns of core taxa summed by relative abundances within bacterial class. “c:” is class and “p:” is phylum.

In contrast to the taxa unique to crops and years, which were rare and not persistent, most of the highly abundant and prevalent taxa were shared (**Figure 4A-C**). We first quantified the abundance and occupancy distributions of OTUs, and then identified OTUs that were consistently detected across replicate plots at one sampling time (occupancy of 1) to include in the core. We found that these 44, 51 and 42 core taxa (as highlighted in **Figure 4A-C**) contribute 84.4%, 79.5% and 79.4% to the total beta diversity in miscanthus 2016, switchgrass 2016 and 2017, respectively (**Figure 4D-F**). While these core taxa were highly abundant and persistent on these crops’ leaves, their functions are yet unknown and additional members could also transiently contribute. However, we suggest that the core taxa identified here should be prioritized for follow-up study of functionality and potential plant benefits.

Notably, if we had defined the core as those taxa uniquely detected on each crop (as in a Venn diagram analysis, **Figure S6**), we would have instead identified rare and transient taxa (**Figure 4A-C**). Thus, a core analysis based on presence and absence, instead of on abundance and occupancy, would have provided a different, and arguably less ecologically relevant, core for these perennial crops. The approach used here provides a reproducible and conservative option for longitudinal series, and allows for systematic discovery ^46,47^ of a replicated core over time.

The core taxa included several Proteobacteria (*Methylobacterium*, *Sphingomonas*, and *Pseudomonas* spp.) and Bacteroidetes (*Hymenobacter* spp.). The taxonomic affiliations of these core taxa are consistent with the literature for other phyllosphere communities ^2,3,18,35,48,49^, providing new support for their seasonal importance in the phyllosphere.

We then performed a hierarchical clustering analysis of standardized (e.g., z-score) dynamics to explore seasonal trends of core taxa (**Figure 4 G-I**, **Figure S7**)^50^. This analysis identified several discrete, seasonally-defined groups of core taxa in switchgrass and miscanthus, respectively. Seasonal groups were taxonomically consistent across crops and years (**Figure 4 J-L**). This finding suggests potential for functional redundancy because closely related taxa are hypothesized to have substantial overlap in their functional repertoire ^51^. The early-season groups (**Figure S7, Figure 4G-I** red traces) included several *Gammaproteobacteria* (**Figure 4J-L**). The late-season groups (**Figure S7**, **Figure 4G-I** blue traces) was comprised of *Alphaproteobacteria* to *Cytophagia* and *Actinobacteria* and was pronounced in switchgrass (**Figure 4J-L**). The third groups **(Figure S7, Figure 4G-I** gray traces) included taxa that peaked in relative abundance mid-season, including *Alphaproteobacteria* and few taxa belonging to *Beta*- and *Gamma*-*proteobacteria*, *Cytophagia, Sphingobacteria* and *Actinobacteria* (**Figure 4J-L**).

Our data suggest a compensatory relationship between members within the Proteobacteria, where members of *Gammaproteobacteria* and *Alphaproteobacteria* replace one another over time (**Figure 5A**). Such community transitions have been observed on the phyllosphere of crops such as sugarcane ^37^, common beans, soybeans, and canola ^38^. A study of endophytic bacteria of prairie grasses, including switchgrass, showed the same trend in abundance of *Gamma*- and *Alphaproteobacteria* ^52^ suggesting that these phyllosphere taxa are facultative endophytes or are similarly affected by the plant development. The benefits plants may gain from these taxa are well characterized (see review from ^53^, however it remains unknown what drives the exclusion of *Pseudomonas* and gives rise to *Alphaproteobacteria* (predominantly *Methylobacteria*) in the phyllopshere and endopshere. One possible explanation would be nutrient availability regulated by the plant development which would selectively influence the abundances of these taxa. Delmotte and colleagues ^54^ hypothesized that *Pseudomonas* specialize on monosaccharides, disaccharides and amino acids, whereas *Sphingomonas* and *Methylobacteria* are generalist scavengers that can subsist on a variety of substrates present at low amounts.

**Figure 5.**
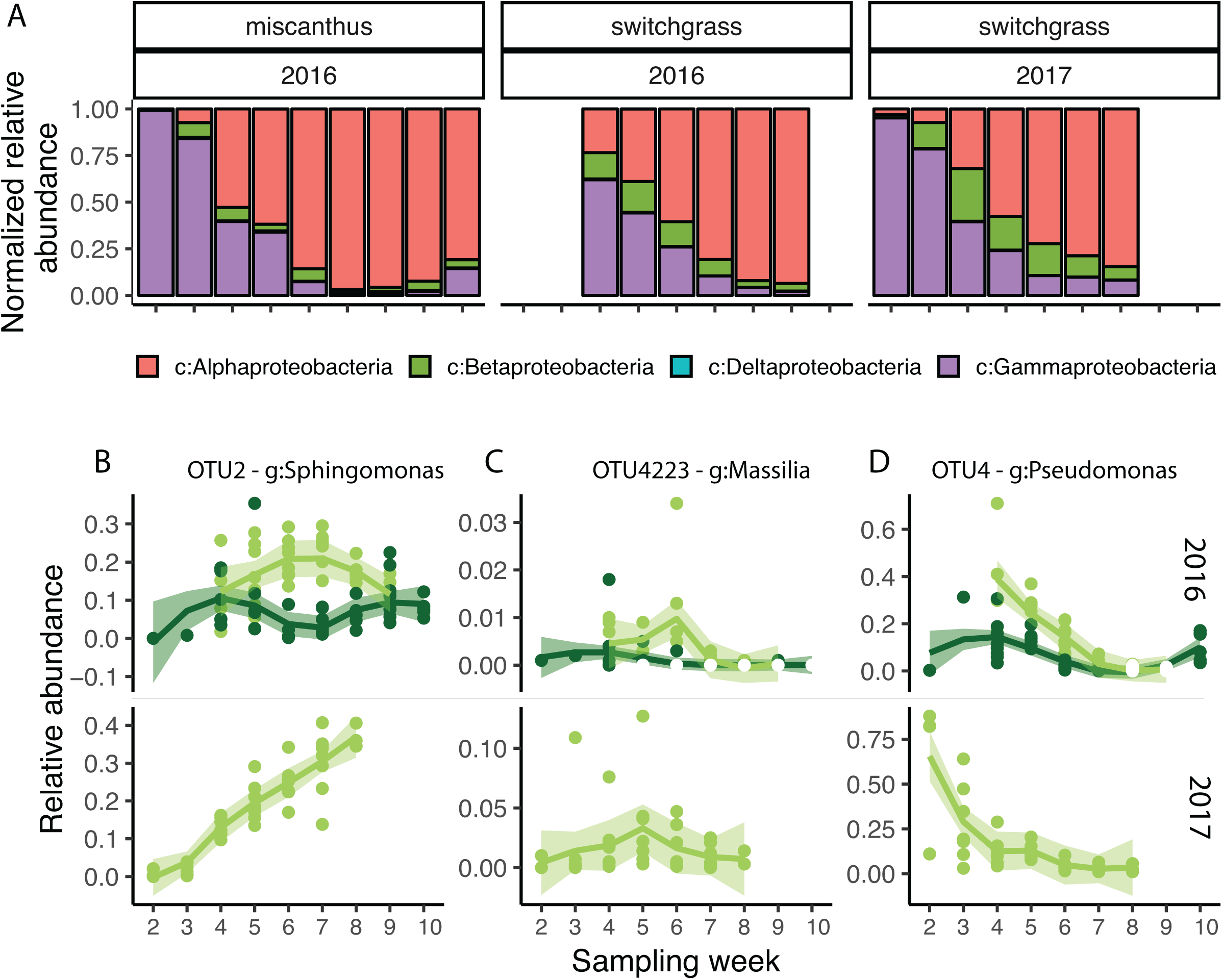
Compensatory patterns of Protobacteria classes over crops and season in the phyllosphere of switchgrass and miscanthus. Proteobacteria OTUs contributed 35.2% of the total taxa detected in the phyllosphere and contributed 116,760 total reads (34.1% of the leaf reads). **(A)** Changes in the relative contributions of all 521 leaf-detected Proteobacteria OTUs by class, over time. **(B-D)** Vignettes showing different dynamics of Proteobacteria OTUs that were detected within the phyllosphere core microbiome, over time and across crops. Class is “c:” and genus is “g:”.

Despite similarity in the membership and dynamics of the core microbiota on both crop plants, there were differences in the relative abundances of the same taxa across crops, suggesting some microbiome adaptation to or selectivity by the host plant (**Figure 5B**). There were other core OTUs that had consistent dynamics across both crops, but these examples demonstrate, first, that dynamics can be crop specific, and second, that abiotic filtering of an OTU to a particular crop could be manifested as differences in dynamics in addition to the more extreme scenarios of taxon exclusion or crop specificity that are the hypotheses posed given unique detection on a particular crop. Indeed, because the crop-unique taxa were generally rare and transient (**Figure 4A-C**), the dynamics of core taxa with crop-distinct dynamics may harbor clues as to the competitive landscape and microbially important changes in the host leaf environment across crops.

### The contributions of abiotic variables, space, time, and crop on phyllosphere assembly

We summarize our analyses of the contributions of crop (host plant), space, time, and abiotic variables to the assembly of the core phyllosphere community in order of least to most important. Spatial distance between the plots had no explanatory value (assessed by distance-decay of beta diversity using a Mantel test with a spatial distance, r: 0.013, p = 0.256). This finding is different from a recent study of annual crops (common beans, canola and soybean) that showed an influenced of sampling location on leaf microbiome structure ^38^. Here, among those that variables were significant, crop (switchgrass or miscanthus) had the lowest explanatory value (**Table 1**). However, our work agrees with previous research that has shown a relationship between plant species/genotype and the leaf microbiota of perennial plants such as wild mustard ^55^, sugar cane ^37^, and tree species like birch, maple, and pine ^56^. Time and measured abiotic factors had highest explanatory value (**Table 1**, **Table S3**). Relatedly, Copeland et al. 2015 showed that stage in plant development can influence leaf microbiome structure in annual crops.

To conclude, we investigated the assembly and seasonal dynamics of the phyllosphere and soil microbes of two perennial grasses, switchgrass and miscanthus, and found consistent community trajectories and memberships across growing seasons, suggesting that their key players are predictable and that most of them can be detected in associated soils. Understanding the seasonal patterns of these key taxa could be used to improve biomass production, plant health, or facilitate conversion. As seen in ^57^, the introduction or control of a few key microbial species can have significant impact on the host plant phenotype. Next steps should be to interrogate core members for functionality and direct interactions with the plant, including investigations of the interactions among core members and with the host crop. This exploration lays the foundation for an approach to biofuel grass production that incorporates an understanding of host-microbe and microbe-microbe interactions.

## Methods

### Site description & sampling scheme

Our study system is located within the Great Lakes Bioenergy Research Center (GLBRC) Biofuel Cropping System Experiment (BCSE) in Hickory Corners, Michigan (42°23’41.6” N, 85°22’23.1” W). We collected samples from two biofuel crops within the BCSE, switchgrass (*Panicum virgatum* L. cultivar “Cave-in-rock”) and miscanthus (*Miscanthus × giganteus)*. Both crops had been continuously grown since 2008, in replicate 30 × 40 m plots arrayed in a randomized complete block design. Within each plot, nitrogen-free (no fertilizer) subplots were maintained in the western-most 3 m of each plot. We sampled replicate plots 1-4 in both the main and the nitrogen free subplots. We collected leaf and bulk soil samples every three weeks across the 2016 growing season, including bare soil in April through senescence in October and November. In total, we collected 152 soil samples (72 switchgrass and 80 miscanthus) and 136 leaf samples (64 switchgrass and 72 miscanthus). At each sampling time, leaves were collected and pooled at three flags along a standardized path within each plot. Leaves were removed from the plant stem using ethanol sterilized gloves, then stored in sterile whirl-pak bags until processing. Bulk soil cores (2 × 10 cm) were collected at the same three locations within a plot, sieved through 4 mm mesh, then pooled and stored in whirl-pak bags. All samples were kept on wet ice for transport, then stored at −80 °C.

Soil physico-chemical characteristics (pH, lime, P, K, Ca, Mg, organic matter, NO_3_-N, NH_4_-N, and percent moisture) were measured by the Michigan State University (MSU) Soil and Plant Nutrient Lab (East Lansing, MI, USA, http://www.spnl.msu.edu/) according to their standard protocols. From each plot, 10 switchgrass leaves or 5 miscanthus leaves were processed for leaf dry matter content according to ^58^. Dried leaves were ground to a fine powder using a Sampletek 200 vial rotator and iron roll bars (Mavco Industries, Lincoln, NE, USA), then carbon and nitrogen were measured on an elemental analyzer (Costech ECS 4010; Costech Analytical Technologies Inc, Valencia, CA, USA). Weather data was collected from the MSU Weather Station Network, for the Kellogg Biological Station location (https://mawn.geo.msu.edu) for each sampling day, and plant height and soil temperature were measured on a per-plot basis.

### Nucleic acid extraction & sequencing

Throughout, we use “microbiome” to refer to the bacterial and archaeal members as able to be assessed with 16S rRNA gene sequence analysis. Soil microbial DNA was extracted using a Powersoil microbial DNA kit (MOBio Inc. Carlsbad, California, USA) according to manufacturer’s instructions. Phyllosphere epiphytic DNA was extracted from intact leaves using a benzyl chloride liquid:liquid extraction, followed by an isopropanol precipitation ^59^, using approximately 5 g of leaves (5-10 switchgrass leaves, or a minimum of 2 miscanthus leaves). Metagenomic DNA from both soil and phyllosphere was quantified using a qubit 2.0 fluorometer (Invitrogen, Carlsbad, CA, USA), and DNA concentrations were normalized between all samples prior to sequencing. Paired-end amplicon sequencing was completed by the Department of Energy’s Joint Genome Institute (JGI) using an Illumina MiSeq sequencer, and using the 16S-V4 (515F-804R) primer set ^60^, according to the JGI’s standard operating protocols, and incorporating chloroplast- and mitochondria-blocking peptide nucleic acids to prevent co-amplification of plastid 16S rRNA gene as described in ^61^.

### Sequence quality control and defining operational taxonomic units

BBDuk (v 37.96) was used to remove contaminants and trim adaptor sequences from reads. Reads containing 1 or more ‘N’ bases, having an average quality score of less than ten or less than 51 bases were removed. Common contaminants were removed with BBMap (v 37.96). Primers were trimmed using cutadapt (v1.17). Reads were merged, dereplicated, clustered into 97% identity with usearch (v10.0.240), and classified against version 123 of the Silva Database ^62^ using sintax ^63^. All reads classified as mitochondria, chloroplast or unclassified were removed before the analysis. Additionally, reads from 4371 OTUs assigned only to the domain level were extracted and reclassified using SINA online aligner (https://www.arb-silva.de/aligner/) ^62^. 696 unclassified reads subsequently were confirmed to be Bacteria could then be classified to more resolved taxonomic levels. Remaining reads were BLASTed against the entire NCBI nucleotide database and specifically against the switchgrass genome to check for non-specific binding, but no hits could be found.

### Alpha and beta diversity

For alpha and beta diversity analyses, we performed analyses to datasets subsampled to the minimum observed quality-filtered reads per sample (141), as well as to 1000 reads per sample. We did this to enable comparison of the most complete time series to the most complete comparative view of diversity. We report richness as total number of OTUs clustered at 97% sequence identity. We used the protest function in the vegan package in R ^64^ to test for synchrony in patterns across crops and years. To calculate beta dispersion, we used the betadisper function in the vegan package in R ^64^, which is a multivariate analogue of Levene’s test for homogeneity of variances. PERMANOVA was used to test hypothesis of beta diversity using adonis function in the vegan package in R ^64^.

### Source-sink models and contributions of soil taxa to leaf communities

Given that virtually all phyllosphere taxa were present in the soil, we evaluated the degree to which observed seasonal patterns in phyllosphere community composition could be explained with a null model. The null model assumed functional equivalence among taxa and random source-sink dynamics from the soil. For each crop, we simulated community dynamics between each pair of sequential samples. This involved: (1) calculating the total number of 0.1 % incremental increases/decreases in OTU relative abundances observed between the two sampling points; (2) randomly and iteratively selecting OTUs, weighting their probability of selection by their relative abundances, to increase or decrease by 0.1 % increments of relative abundance until reaching the total number of observed increases/decreases for the sample pair; (3) counting the number of immigrant OTUs which appeared only in the second sample of the pair; (4) randomly selecting OTUs in the simulated community to decrease by 0.1 % increments until the total decrease equaled the observed number of arriving OTUs multiplied by their median initial abundance (0.1 %); and then (5) randomly selecting, again weighting their probability of selection by their relative abundance, the predicted number of immigrant OTUs from the soil community, such that the final simulated community abundance was equal to 1000 sequences. The source soil community was generated by pooling all soil samples from both years. We used this process to simulate demographic stochasticity. In the model, it was assumed that 0.1 % incremental changes in relative abundance realistically reflects phyllosphere population dynamics, and that an immigration event results in the initial relative abundance of an OTU at 0.1%. Importantly, even if these assumptions are imprecise, the simulation provides a consistent baseline of community composition against which to compare observations over the growing season.

### Core taxa selection

To infer the core phyllosphere taxa and prioritize them for further inquiry, we calculated the abundance-occupancy distributions of taxa, as established in macroecology (e.g. Shade et al. 2018). For each OTU, we calculated occupancy and mean relative abundance at each time point by crop and year. Only OTUs with occupancy of 100% (found in all samples at a particular time point) were prioritized as core members. Using this conservative threshold for occupancy, we included all OTUs that had strong temporal signatures; these taxa also were in high abundance and were persistent as indicated by their abundance-occupancy distributions. These core taxa also represent potentially important players in plant development, as they were detected at least at one time point in all sampled fields.

We quantified the explanatory value of the core members to community temporal dynamics using a previously published method of partitioning community dissimilarity ^65^:

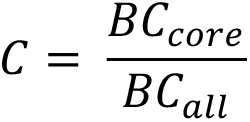

Where *C* is the relative contribution of community Bray Curtis (*BC*) dissimilarity attributed to the core OTUs.

### Hierarchical clustering

To understand the seasonal abundance patterns of the core taxa we performed hierarchical clustering. We used a z-scored relative abundance matrix subset to contain only core taxa to generate a w complete linkage distance matrix using the R function hclust() ^50^. Groups of core taxa with similar dynamics were defined from the dendrogram using the function cutree() in R with number of desired groups (k=) to be close to the number of sampling time points; 8 for miscanthus 2016, 5 for switchgrass 2016 and 7 for switchgrass 2017.

### Availability of data, workflows, and material

The datasets generated and/or analyzed during the current study are available in the Joint Genomes Institute, Integrated Microbial Genomes repository with JGI Projects designated by year and sample type (Project ID 1139694, 1139696 for 2016 season phyllosphere and soil, and 1191516 and 1191517 for 2017 season phyllosphere and soil sequences, respectively). Our sequence analyses and statistical workflows are available at https://github.com/ShadeLab/PAPER_GradySorensenStopnisek_InPrep

## Authors’ contributions

AS designed the study. AS, KLG, JS, and NS conducted field work. KLG executed lab work. JS, NS, JG, and AS analyzed the data. All authors discussed and revised the manuscript.

## Acknowledgements

We thank SH Lee, M Sleda, S Wu and M Nunez for technical assistance in the field and laboratory. This material is based upon work supported by the Great Lakes Bioenergy Research Center, U.S. Department of Energy, Office of Science, Office of Biological and Environmental Research under Award Numbers DE-SC0018409 and DE-FC02-07ER64494. The work conducted by the U.S. Department of Energy Joint Genome Institute, a DOE Office of Science User Facility, is supported under Contract No. DE-AC02-05CH11231. This work was supported in part by Michigan State University through computational resources provided by the Institute for Cyber-Enabled Research. NS acknowledges support from the Michigan State Plant Resilience Institute.

## Supporting Figures

**Figure S1.**
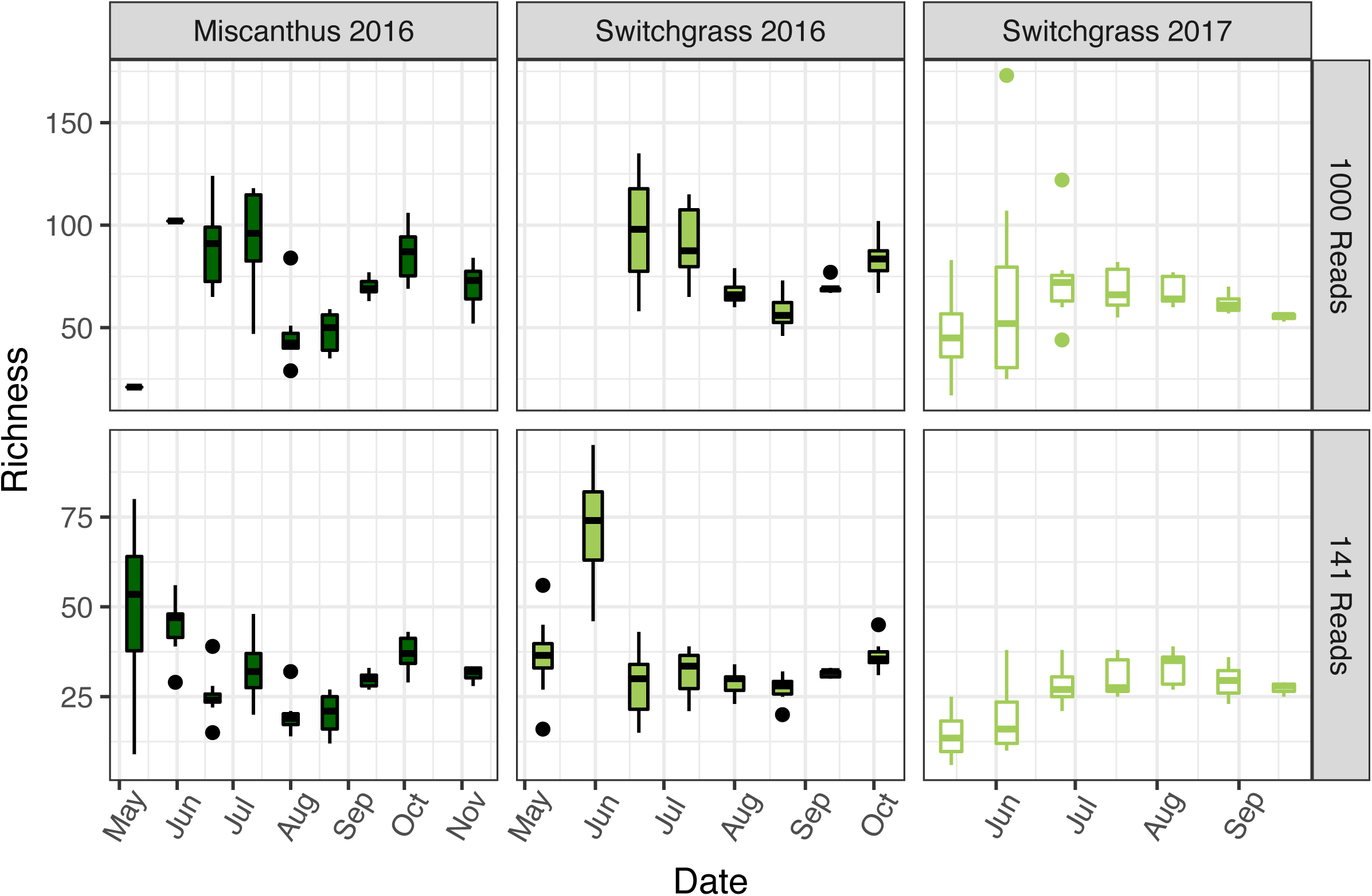
Seasonal patterns in the number of observed phyllosphere taxa (richness). Operational taxonomic units (OTUs) were defined at 97% amplicon sequence identity. Richness is provided at subsampling depths of 1000 reads (top) and 141 reads (bottom).

**Figure S2.**
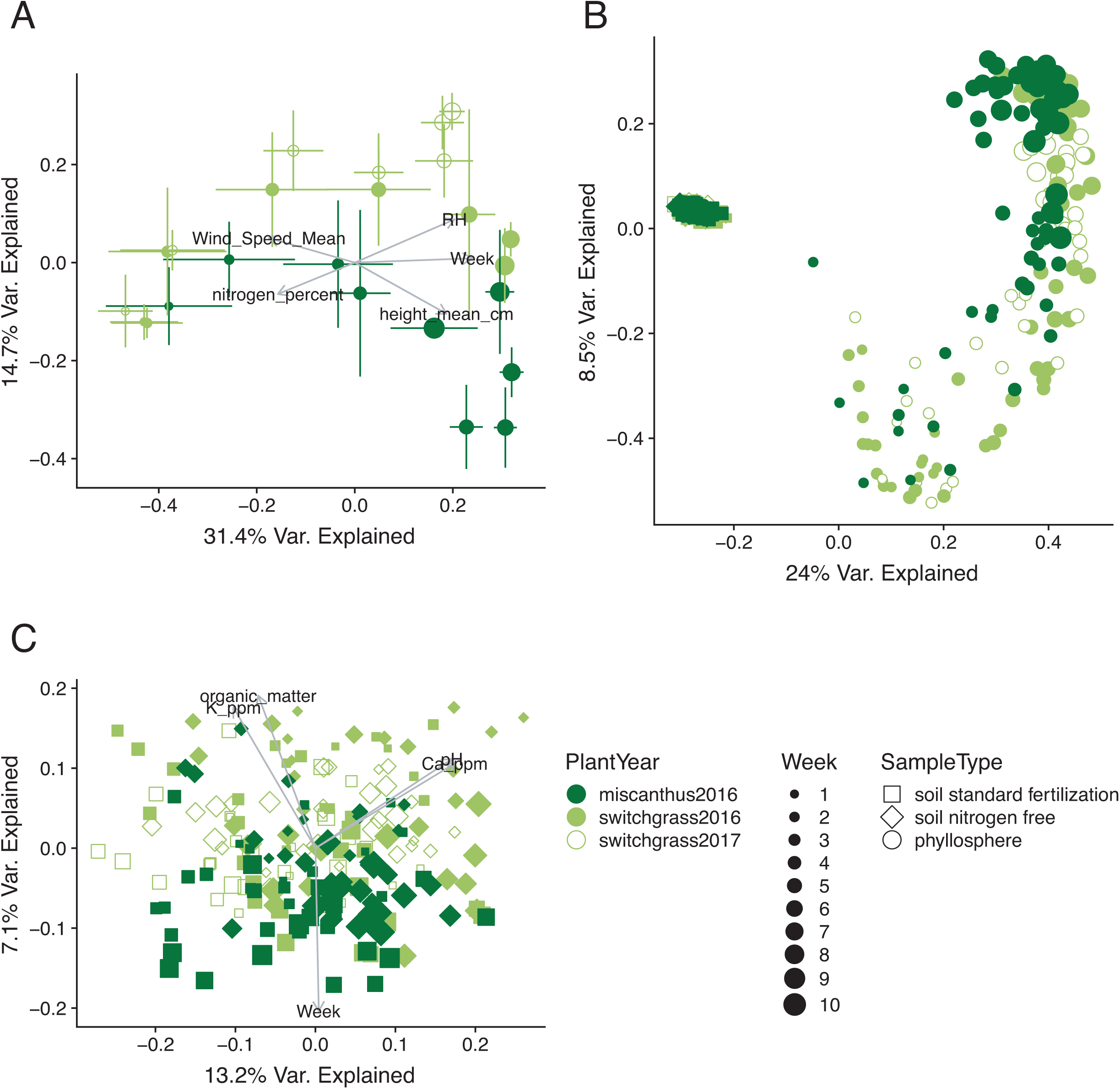
Seasonal patterns in the structures of bacterial and archaeal communities inhabiting the phyllosphere and associated soils of the biofuel feedstocks switchgrass and miscanthus. **(A)** Principal coordinates analysis (PCoA) of switchgrass and miscanthus phyllosphere communities (Bray-Curtis dissimilarity), error bars show 1 deviation around the centroid (n = 3 to 8 replicate plots/time point). Subsampling depth was 141 reads per sample and environmental vectors are fitted when r^2^ > 0.4 and p < 0.05. **(B)** PCoA of the phyllosphere communities relative to the soil, subsampled to 141 sequences per sample. **(C)** PCoA of the soil communities associated with miscanthus and switchgrass, subsampled to 19,967 sequences per sample.

**Figure S3.**
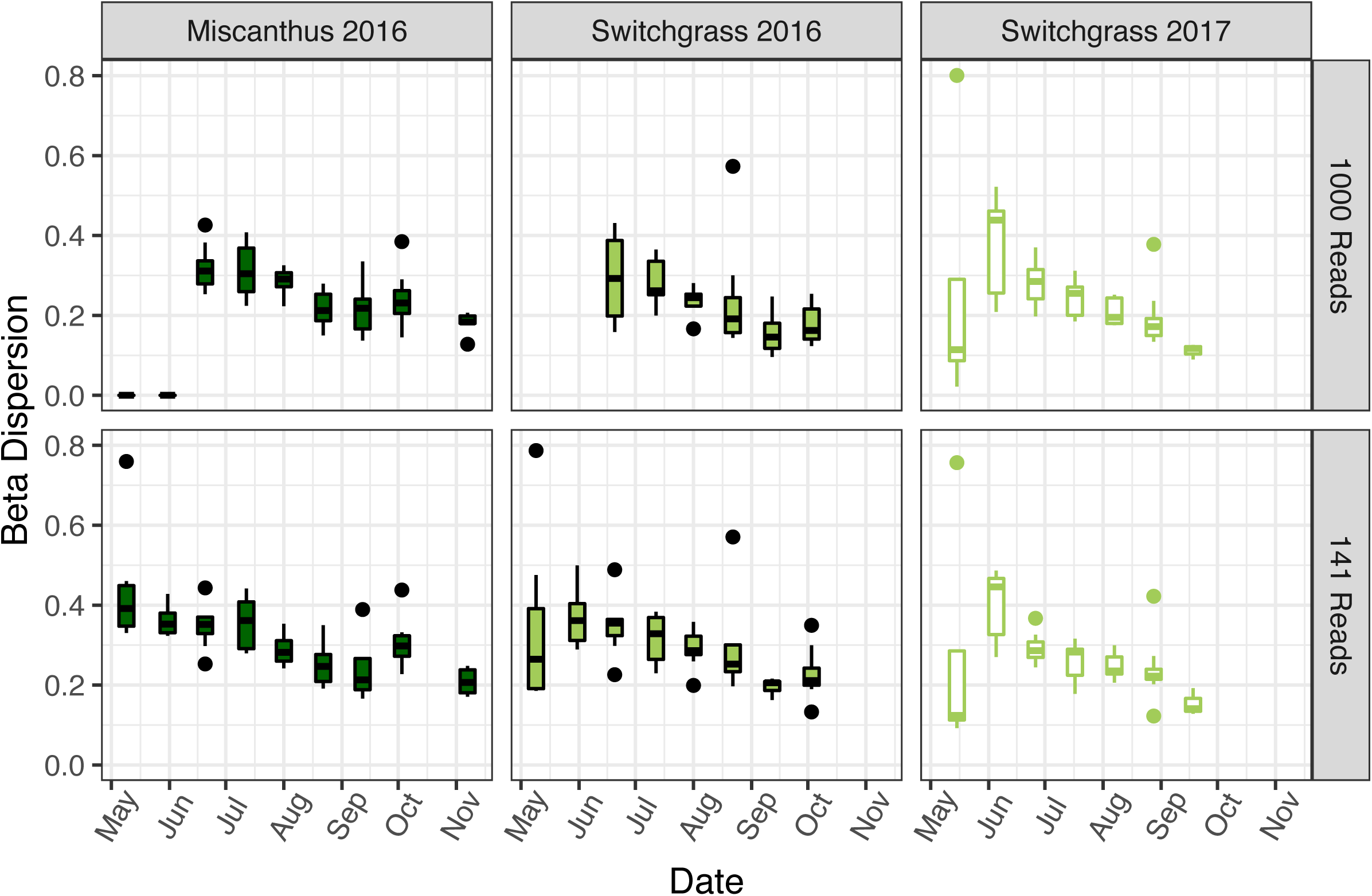
Phyllosphere communities become less variable over time. Distance to median was calculated by analysis of beta-dispersion. Variability in phyllosphere microbiome structure over time miscanthus 2016, switchgrass 2016, and switchgrass 2017 field seasons. Betadispersion was calculated from data series subsampled to 1000 reads (top) and 141 reads (bottom).

**Figure S4.**
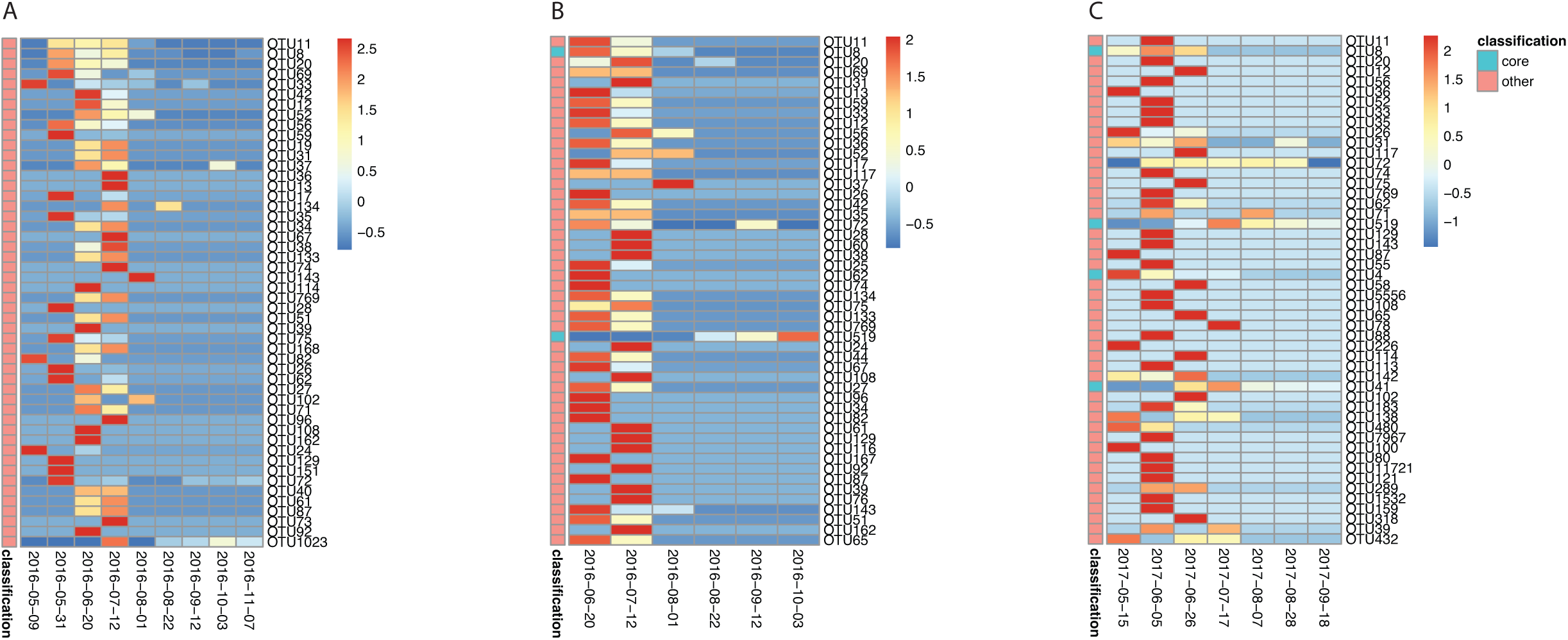
Decreases in the contributions of soil-dominating taxa to the phyllosphere microbiome over time. Heatmaps represent the 50 top-ranked OTUs from the soil that were also detected in the phyllosphere. The cell colors are the z-scored relative abundances of the OTUs in the phyllosphere. The ranking is from top to bottom (e.g., the most abundant soil-dominant taxon that was also detected in the phyllosphere is represented by the top row in the heatmap). The left bar shows the classification the taxon as either a core member (green) or not (pink). **(A)** Miscanthus 2016; **(B)** switchgrass 2016; **(C)** switchgrass 2017.

**Figure S5.**
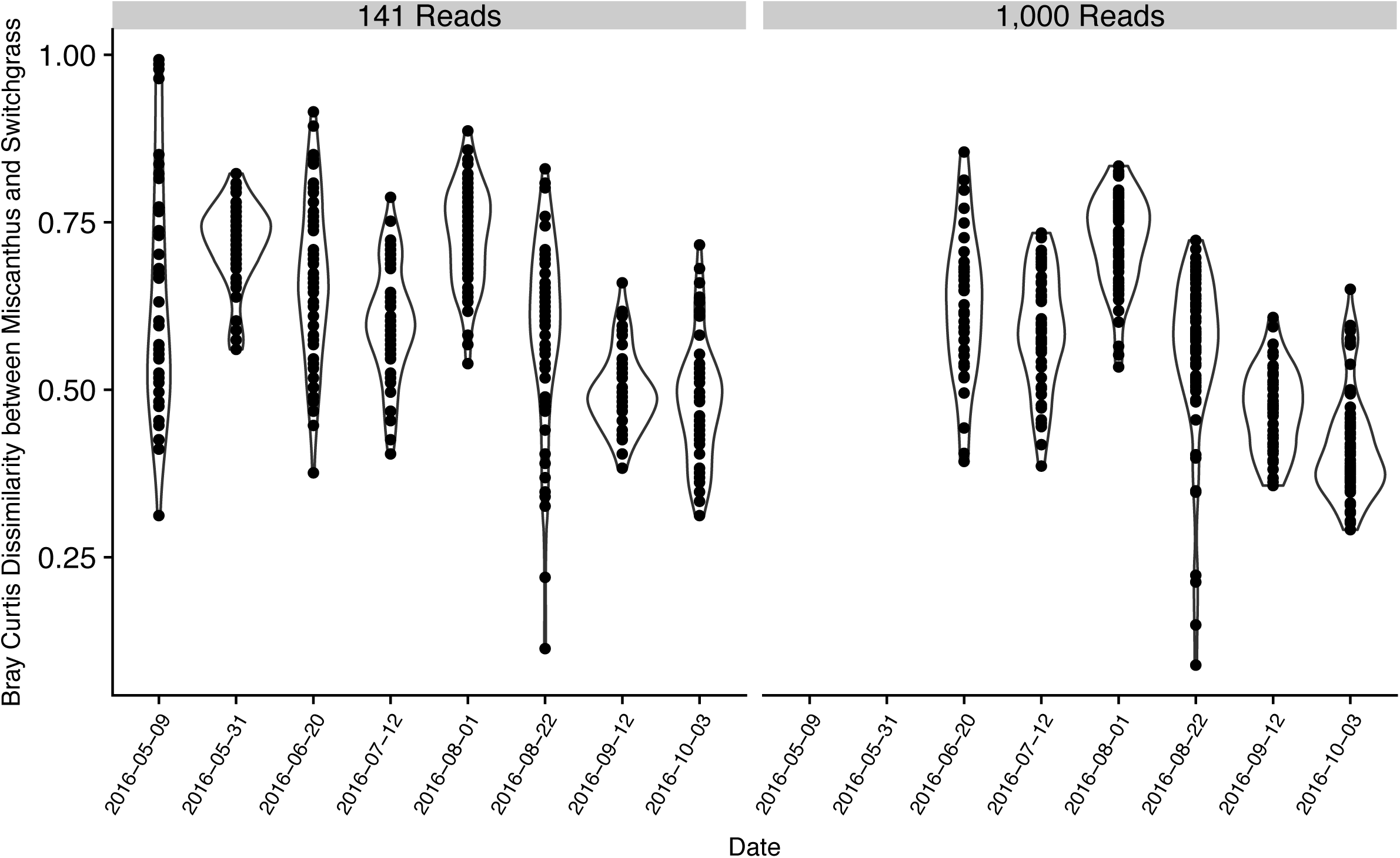
Intra-crop similarity in leaf communities over time in 2016. We show changes in Bray-Curtis dissimilarities between switchgrass and miscanthus phyllosphere microbiomes per time point, inclusive of the maximum number of replicated blocks (up to 8) per time point. **(A)** “Full” time series, subsampled to 141 reads per sample. **(B)** Time series subsampled to 1000 reads per sample.

**Figure S6.**
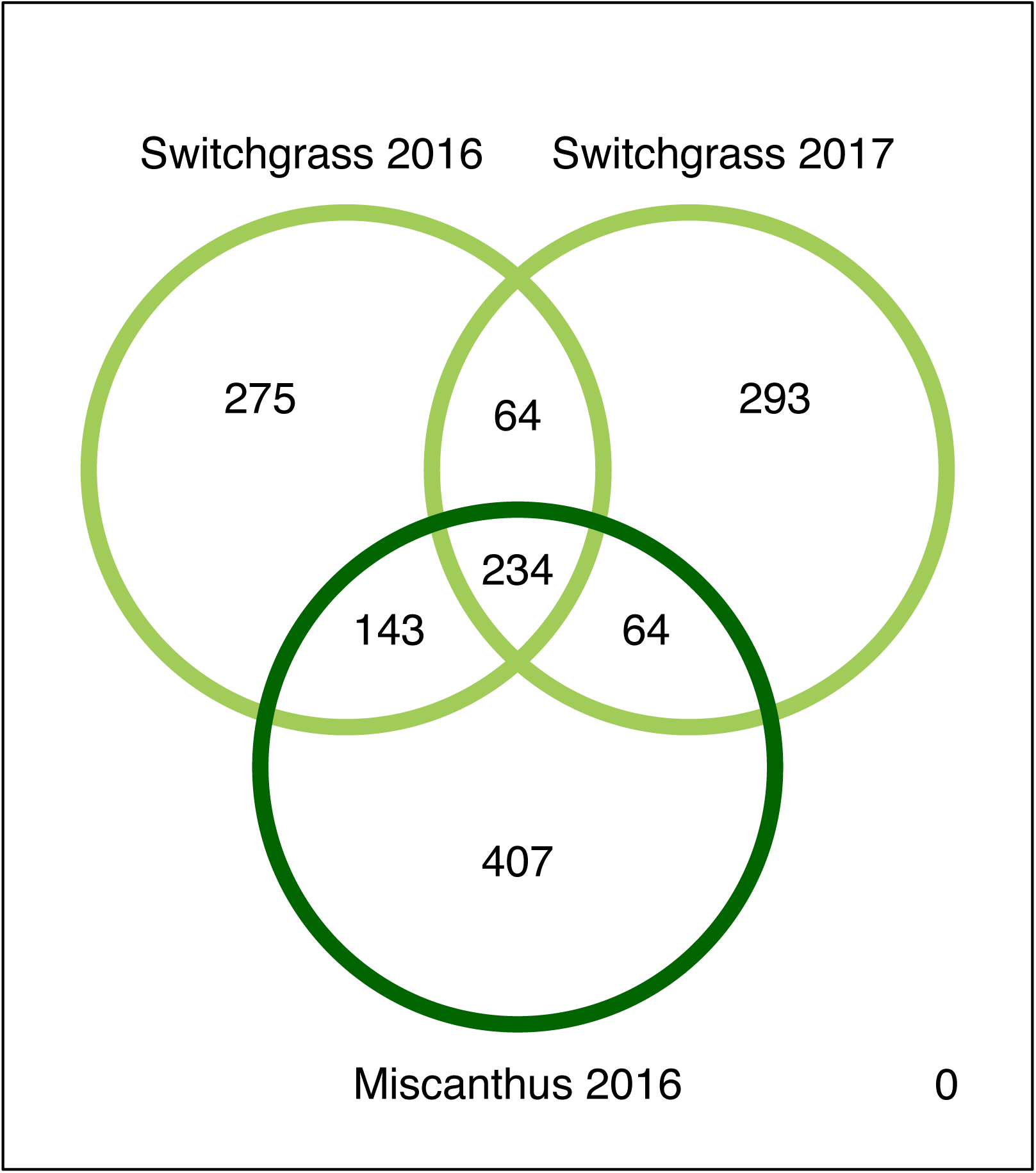
Venn diagram of taxa shared across the switchgrass and miscanthus phyllosphere in 2016 and 2017. Data were rarefied to 1000 reads per sample.

**Figure S7.**
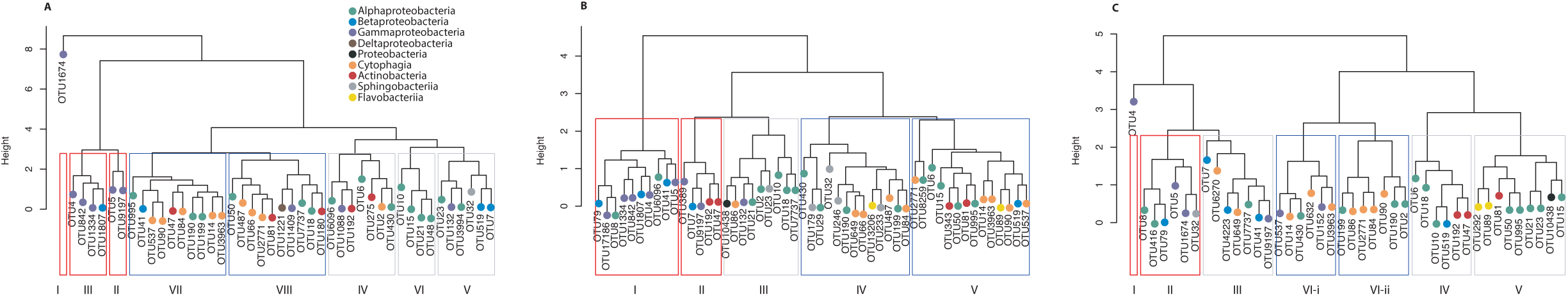
Hierarchical clustering of standardized (z-scored) dynamics of core phyllosphere taxa on the phyllosphere of miscanthus 2016 (A), switchgrass 2016 (B) and switchgrass 2017 (C). Seasonally discrete clusters coincide with plant phenology, including groups that achieved highest relative abundance during early (red), mid (gray) and late (blue) plant growth. Clusters are labeled in order of temporal occurrence in **Figure 4G-I**. Circles on the dendrogram tips are color coded by OTU taxonomic classification and labeled with the OTU ID.

## Supporting Tables

**Table S1.** **Sequencing summary of phyllosphere microbial communities characterized in this study,** categorized by crop and year.

|  | Miscanthus<br>2016 | Switchgrass<br>2016 | Switchgrass<br>2017 |
| --- | --- | --- | --- |
| Raw Read Pairs | 7336923 | 7120682 | 9271169 |
| QC Reads | 7142900 | 6883631 | 7123783 |
| % Chloroplast/ Mitochondria of QC Reads | 77.27% | 67.16% | 12.32% |
| Samples >= 141 Reads | 65 | 62 | 44 |
| Samples >= 1,000 Reads | 53 | 43 | 44 |
| Samples >= 10,000 Reads | 36 | 27 | 44 |
| OTUs (rarefied to 141 Reads) | 540 | 685 | 232 |
| OTUs (rarefied to 1,000 Reads) | 1010 | 859 | 769 |
| OTUs (rarefied to 10,000 Reads) | 1121 | 896 | 2320 |

**Table S2.** Comparison of overarching patterns of beta diversity across the same dataset rarefied to different sequencing depths. We compared all pairs of 141, 500, 1000, 5000, or 10000 reads per sample. All Mantel tests were significant at p < 0.001 on 1000 permutations.

|  | 141 Reads | 500 Reads | 1000 Reads | 5000 Reads |
| --- | --- | --- | --- | --- |
| 500 Reads | 0.932 |  |  |  |
| 1000 Reads | 0.872 | 0.982 |  |  |
| 5000 Reads | 0.752 | 0.931 | 0.976 |  |
| 10000 Reads | 0.725 | 0.916 | 0.967 | 0.999 |

**Table S3.** Fitted environmental variables that explain changes in microbiome community structure. Values in which EnvFit R^2^ > 0.40 were plotted as vectors in **Figure 2**.

| Variable Tested | Axis 1 | Axis 2 | R squared | P -value |
| --- | --- | --- | --- | --- |
| precipitation | -0.127 | 0.146 | 0.038 | 0.703 |
| Air_temp_mean | -0.298 | -0.078 | 0.095 | 0.384 |
| air_temp_max | -0.212 | -0.056 | 0.048 | 0.633 |
| Air_Temp_Min | -0.242 | 0.139 | 0.078 | 0.439 |
| Air_Pressure | -0.022 | 0.217 | 0.048 | 0.653 |
| RH | 0.474 | 0.444 | 0.422 | 0.008 |
| AH | -0.081 | 0.102 | 0.017 | 0.834 |
| Wind_Speed_Mean | -0.703 | -0.213 | 0.539 | 0.001 |
| Solar_Radiation | -0.258 | -0.368 | 0.202 | 0.118 |
| PAR | -0.300 | -0.288 | 0.173 | 0.162 |
| soil_temp_5_cm_bare_avg | -0.408 | -0.061 | 0.170 | 0.163 |
| Week | 0.730 | 0.356 | 0.659 | 0.001 |
| LDMC_mg_per_g | 0.418 | 0.177 | 0.206 | 0.001 |
| nitrogen_percent | -0.392 | -0.416 | 0.327 | 0.001 |
| carbon_percent | 0.345 | 0.038 | 0.121 | 0.001 |
| carbon_per_nitrogen | 0.274 | 0.134 | 0.093 | 0.003 |
| height_mean_cm | 0.730 | -0.230 | 0.586 | 0.001 |
| pH | -0.027 | 0.219 | 0.049 | 0.047 |
| P_ppm | -0.120 | -0.097 | 0.024 | 0.187 |
| K_ppm | -0.445 | 0.187 | 0.233 | 0.001 |
| Ca_ppm | 0.047 | 0.059 | 0.006 | 0.659 |
| Mg_ppm | -0.366 | -0.125 | 0.149 | 0.001 |
| organic_matter | -0.348 | -0.036 | 0.123 | 0.001 |
| NO3N_ppm | 0.003 | -0.080 | 0.006 | 0.646 |
| NH4_ppm | -0.461 | 0.090 | 0.220 | 0.001 |
| soil_moisture_percent | 0.609 | 0.015 | 0.371 | 0.001 |
| soil_temp_10cm | -0.364 | -0.028 | 0.133 | 0.001 |

## Datasets

**Dataset 1.** Operational taxonomic unit identifiers and representative 16S rRNA gene amplicon sequences of the core switchgrass and miscanthus phyllosphere microbiota.

